## Supplementary Data for "Targeting endometrial cancer stem cell activity with metformin is inhibited by patient-derived adipocyte-secreted factors"

Supplementary Tables

S1. Taqman assays used for gene expression analysis.

| Gene symbol | Gene Name | Assay ID |
|---|---|---|
| SOX2 | *SOX2* | Hs01053049_s1 |
| NANOG | *NANOG* | Hs02387400_g1 |
| BMI1 | *BMI1* | Hs00995520_g1 |
| HEY1 | *HEY1* | Hs01114113_m1 |
| HES1 | *HES1* | Hs00172878_m1 |
| WNT2 | *WNT2* | Hs00608224_m1 |
| CTNNB1 | *B-CATENIN* | Hs00355045_m1 |
| YAP1 | *YAP1* | Hs00902712_g1 |
| TWIST1 | *TWIST* | Hs01675818_s1 |
| SNAI1 | *SNAI1* | Hs00195591_m1 |
| ZEB1 | *ZEB1* | Hs01566408_m1 |
| CDH1 | *E-CADHERIN* | Hs01023895_m1 |
| OCLN | *OCCLUDIN* | Hs00170162_m1 |
| CLDN3 | *CLAUDIN* | Hs00265816_s1 |
| DSP | *DESMOPLAKIN* | Hs00950591_m1 |
| EPCAM | *EPCAM* | Hs00901885_m1 |
| VIM | *VIMENTIN* | Hs00958111_m1 |
| CDH2 | *N-CADHERIN* | Hs00983056_m1 |
| PGK1 | *PGK1* | Hs00943178_g1 |

S2. Characteristics of patients from whom omental biopsies were obtained for primary pre-adipocyte culture and maturation.

| **Patient ID** | **Age** | **BMI** | **Presence of diabetes** | **Grade of endometrial cancer** | **FIGO stage of endometrial cancer** | **Trial treatment allocation** |
|---|---|---|---|---|---|---|
| C21 | 59 | 54.2 | No | 1 | 1a | Metformin |
| O15 | 53 | 36.6 | No | 2 | 3a | Metformin |
| S40 | 51 | 25.3 | No | 3 | 2 | Placebo |
| W06 | 64 | 29.4 | No | 1 | 1a | Metformin |
| S33 | 61 | 41.4 | No | 1 | 2 | Placebo |
| C27 | 78 | 28.5 | No | 3 | 1b | Placebo |

Supplementary Figures

S1. Relationship between $ALDH^{high}$ and $CD133^{+ve}$ cells in Ishikawa and Hec-1a cell lines (n=5).

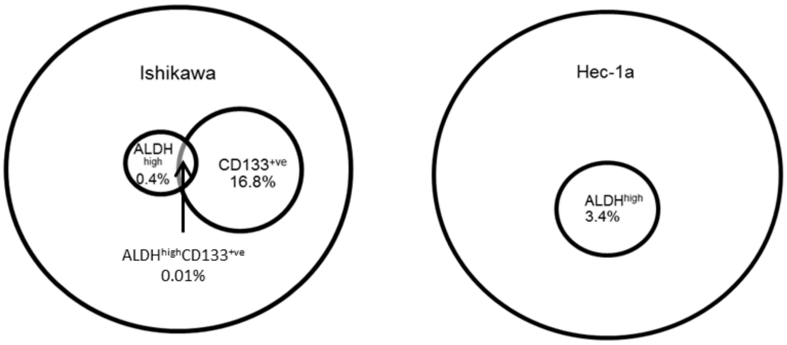

S2. Expression of genes associated with a cancer stem cell phenotype in ALDH$^{high}$ and CD133$^{+ve}$ endometrial cancer cells. **(a)** RT-qPCR of genes associated with pluripotency and self-renewal in ALDH$^{high}$ and CD133$^{+ve}$ cells (n=3). **(b)** RT-qPCR of EMT transcription factor genes in ALDH$^{high}$ and CD133$^{+ve}$ cells (n=3).

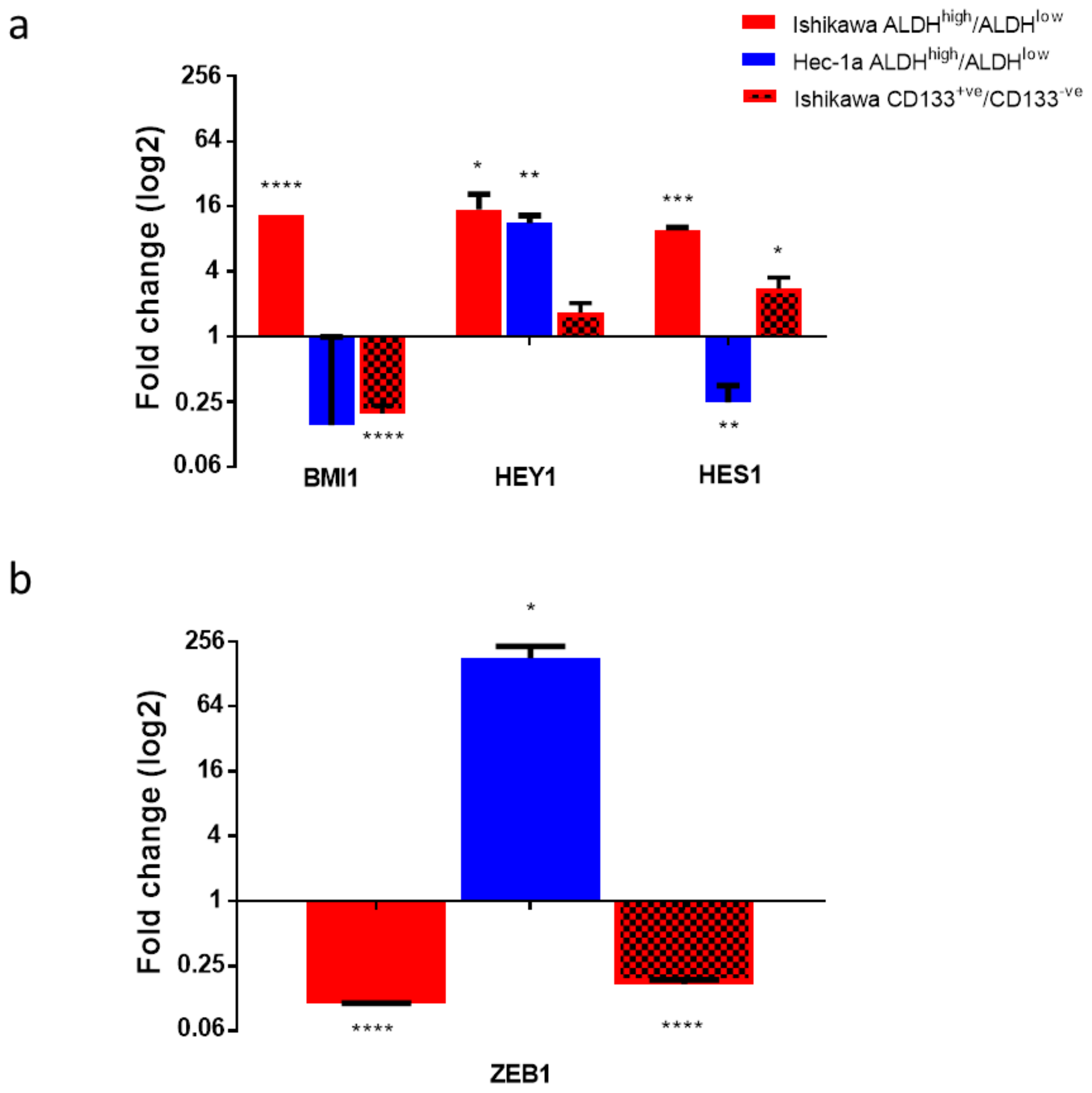

S3. Effect of metformin on endometrial cancer stem cell number and activity. **(a)** Self-renewal of sphere-initiating Ishikawa and Hec-1a cells treated with metformin in the first generation (n=4). **(b)** Sulforhodamine B (SRB) cytotoxicity assay of Ishikawa and Hec-1a cells treated with metformin (n=3).

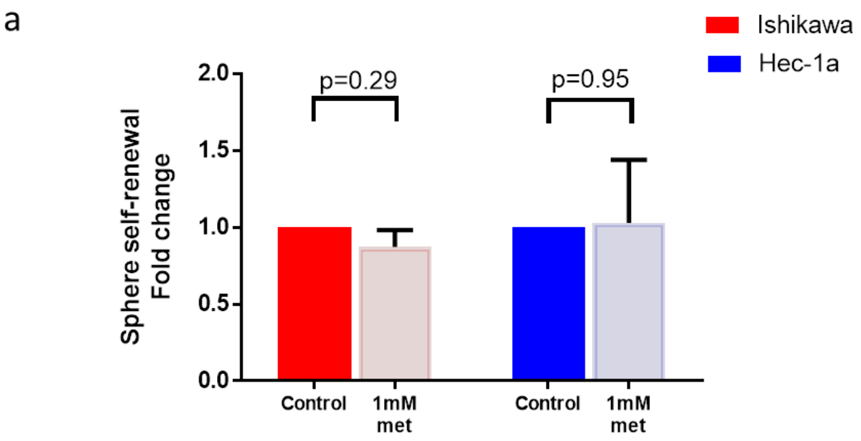


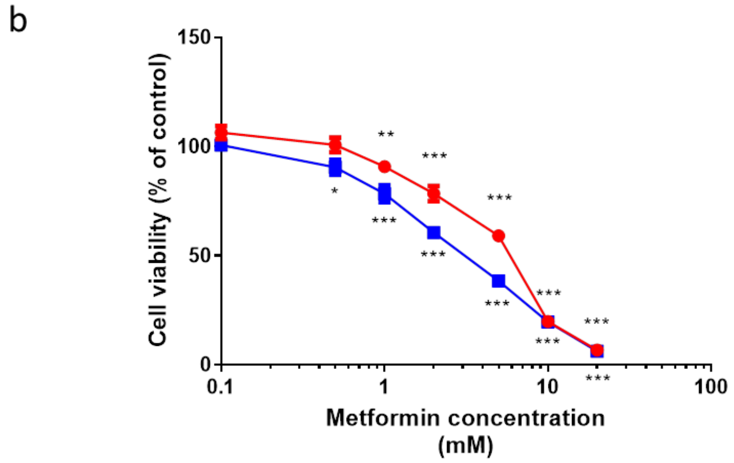

S4. Effect of metformin on stem cell and EMT gene expression. **(a)** On the top, qRT-PCR of genes within the Wnt and Hippo signalling pathways in Ishikawa cells treated with metformin. Underneath, qRT-PCR of the same genes in Hec-1a cells treated with metformin (n=3). **(b)** On the top, qRT-PCR of EMT transcription factor genes in Ishikawa cells treated with metformin. Underneath, qRT-PCR of the same genes in Hec-1a cells treated with metformin (n=3). Data are represented as the mean±SEM. *p≤0.05, **p≤0.01, ***p≤0.001, ****p≤0.0001.

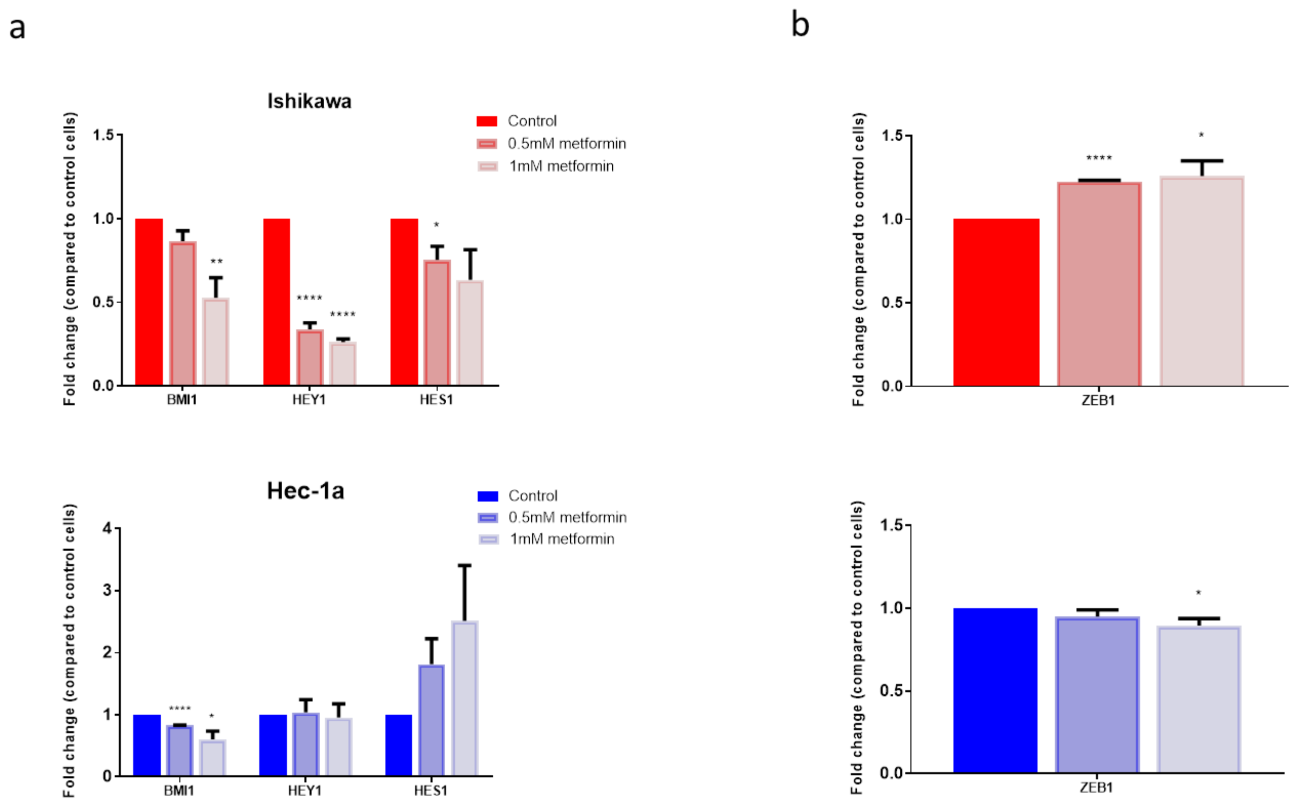

S5. Effect of pre-adipocyte conditioned media (PACM) on endometrial cancer stem cell activity and response to metformin. **(a)** images at x100 magnification of pre-adipocytes exposed to control growth media for 15 days before being stained with Oil Red O. **(b)** SRB cytotoxicity assay of Ishikawa cells exposed to PACM and treated with metformin (n=4). **(c)** SFE of Ishikawa cells treated with PACM (n=6).

a

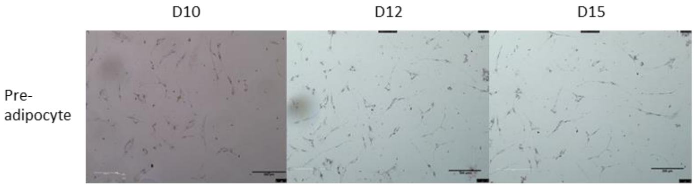


b

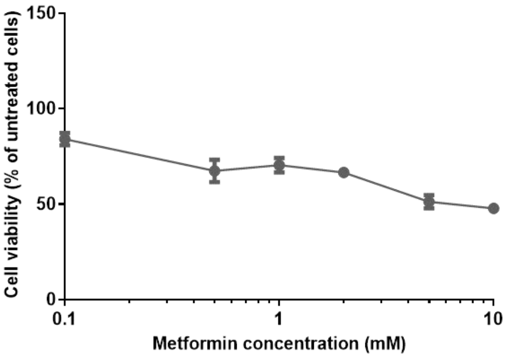


c

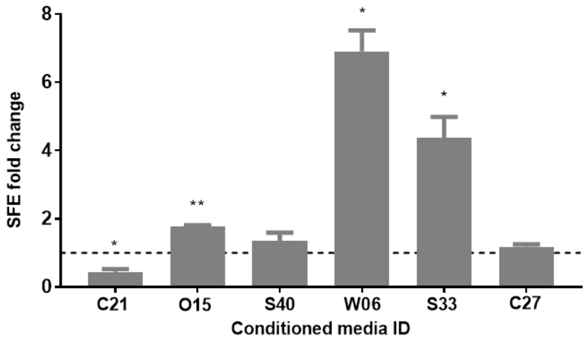

S6. *In vivo* effect of metformin on ALDH and CD133 expression in endometrial tumour biopsies. **(a)** Baseline characteristics of participants in pre-surgical window study. **(b)** Effect of metformin and placebo on immunohistochemical expression of ALDH (upper) and CD133 (lower) (n=67).

a

| Characteristic | Metformin arm (n=33) | Placebo arm (n=34) |
|---|---|---|
| Age, years | 65.1 (33.3-83.7) | 67.6 (39.8-80.8) |
| BMI, kg/m$^2$ | 32.1 (20.2-54.2) | 32.0 (17.8-47.6) |
| Tumour grade | | |
| AEH | 0 (0.0) | 1 (2.9) |
| 1 | 18 (54.5) | 17 (50.0) |
| 2 | 10 (30.3) | 11 (32.4) |
| 3 | 4 (12.1) | 5 (14.7) |
| 4 | 1 (3.0) | 0 (0.0) |
| FIGO stage | | |
| 1a | 22 (66.7) | 17 (50.0) |
| 1b | 4 (12.1) | 9 (26.5) |
| 2 | 0 (0.0) | 4 (11.8) |
| 3 | 7 (21.2) | 3 (8.8) |
| 4 | 0 (0.0) | 0 (0.0) |
| n/a | 0 (0.0) | 1 (2.9) |

b

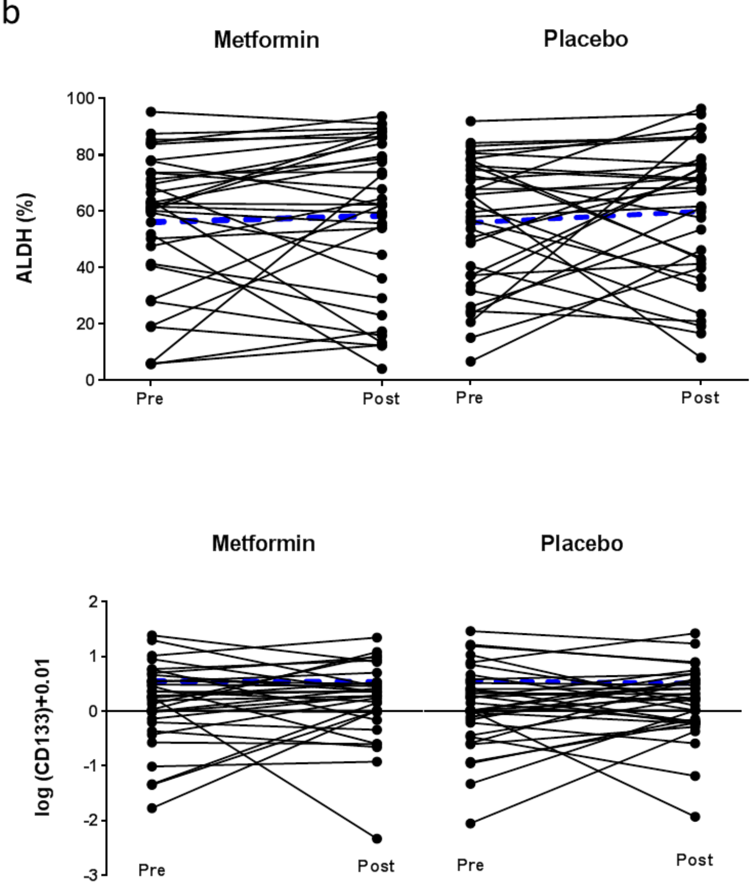
