## Supplementary methods for "Targeting endometrial cancer stem cell activity with metformin is inhibited by patient-derived adipocyte-secreted factors"

Endometrial cancer cell lines

Ishikawa cells represent a well differentiated cell line, expressing oestrogen and progesterone receptors, and have wild-type K-Ras and were purchased from HPA Culture Collection (Salisbury, UK)(1). Hec-1a is a moderately differentiated cell line with defective oestrogen receptors, activating K-Ras mutations and positive PTEN expression and was obtained from ATCC (Middlesex, UK)(1).

Adipocyte culture

The omental tissue was washed twice in phosphate buffered saline (PBS) to remove excess blood. Tissue was finely dissected into 1-2mm^3^ pieces and placed in a centrifuge tube containing collagenase/dispase dissolved in pre-adipocyte growth media to give a final concentration of 1mg/ml. The tissue suspension was placed on a flask shaker for two hours at 37^o^C to aid digestion before being filtered through a 100µm filter to remove connective tissue and undigested fat. Following centrifugation, the cell pellet was resuspended in pre-adipocyte growth media and transferred to a humidified incubator for culture.

Oil Red O staining was used to detect intracellular lipids as confirmation of attainment of a mature adipocyte phenotype. A working solution of three parts Oil Red O stock solution (Sigma-Aldrich, Dorset, UK) diluted with two parts distilled water was made and filtered through a 0.2µm syringe filter to remove any debris. Growth media was aspirated from wells containing confluent mature adipocytes and 10% formaldehyde in PBS was added and left for 10 minutes for fixation to occur. Cells were subsequently washed twice with PBS before the working solution of Oil Red O was applied and left for 15 minutes. Residual extracellular stain was removed by repeated washing with distilled water before cells were visualised under a light microscope.

Sphere formation and passaging

A single cell suspension was created by manual disaggregation (25 gauge needle) and either 5000 (Ishikawa) or 2000 (Hec-1a) cells were seeded in each well of a poly-HEMA (poly (2-hydroxyethylmethacrylate)) coated six well plate in stem cell media (phenol free, high glucose DMEM/F12 [Gibco, Paisley, UK] with 1% penicillin/streptomycin, 20ng/ml EGF and 10ml B27 [Gibco, Paisley, UK]) in the presence or absence of metformin (Sigma-Aldrich, Dorset, UK). Cells were cultured for five days before spheres with a diameter >50µm were counted at x40 magnification using a light microscope. Sphere formation efficiency (SFE) was calculated by dividing the number of spheres formed by the number of single cells initially plated. To assess self-renewal, spheres were counted, mechanically disrupted, centrifuged (450g) and dissociated through the addition of trypsin (Sigma-Aldrich, Dorset, UK) for 90 seconds. Single cells were re-plated at the original seeding density. The self-renewal capacity was determined by dividing the number of secondary spheres formed after five days in culture by the total number of primary spheres.

Sulforhodamine B (SRB) cytotoxicity assay

Cells were plated at a density of 1000 cells/well or 10,000 cells/well (in the case of conditioned media experiments where cell proliferation was markedly reduced due to the omission of FBS from the growth media). After 24 hours, growth media was either replaced with fresh growth or conditioned media in the presence or absence of metformin and maintained in culture for four (conditioned media experiments) or five days. Cells were then fixed with 10% trichloracetic acid for one hour at 4-8^o^C before being washed and left to dry overnight. The following day, 0.4% SRB (Sigma-Aldrich, Dorset, UK) was added and left for 15 minutes before unbound stain was removed by washing with 1% acetic acid. Bound protein was solubilised with 10mM Tris and absorption measured.

Real Time quantitative-Reverse Transcriptase-Polymerase Chain Reaction (RT-qPCR)

Quantification and quality assurance of extracted RNA was performed using the Agilent 2100 Bioanalyzer and Qubit RNA HS Assay Kit (Fisher Scientific, Loughborough, UK).

Reverse transcription of extracted RNA was performed on a thermal cycler (PTC-200 Thermal Cycler, MJ Research, Minnesota, USA) for 5 minutes at 25^o^C, 30 minutes at 42^o^C and 5 minutes at 80^o^C.

Conditions for pre-amplification were 10 minutes at 95^o^C, 15 cycles of 15 seconds at 95^o^C and 4 minutes at 60^o^C. After cycling, the reaction was diluted 1:5 using DNA suspension buffer (Teknova, Hollister, USA) and stored at -20^o^C until required.

Assay stocks (x10) for the RT-qpCR were prepared using the Taqman gene expression assays of interest and a house keeper gene, phosphoglycerate kinase 1 (PGK1), diluted in Assay Loading Reagent (PN 100-5359, Fluidigm, San Francisco, UK). RT-qPCR was performed using a thermal mix consisting of 30 minutes at 25^o^C, 60 minutes at 70^o^C and 2 minutes at 50^o^C followed by a Hot Start of 1 minute at 95^o^C and 35 cycles of 5 seconds at 96^o^C and 20 seconds at 60^o^C.

Fold change was calculated using the formula:

fold change=2-∆∆Ct

where Ct is the threshold cycle, ∆Ct=Ct of the specific gene-Ct of the housekeeper gene PGK1 and ∆∆Ct=∆Ct of ALDH^high^/CD133^+ve^ cells-∆Ct of ALDH^low^/CD133^-ve^ cells

Immunohistochemistry

Sufficient tumour was available for immunohistochemistry to be performed on 34/43 receiving placebo and 33/45 women receiving metformin. Staining of 4µm sections was performed using the Leica Bond Max with heat induced epitope retrieval at pH 6 (CD133) or pH 9 (ALDH). Primary antibody incubation was performed for one hour at room temperature with CD133 (#130-090-422, Miltenyi Biotec, Surrey, UK, dilution 1:25) or ALDH (611194, BD Biosciences, Oxford, UK, dilution 1:100).

Slides were digitised with the Leica SCN400 slide scanner. The percentage of cells with positive apical membrane (CD133) or cytoplasmic (ALDH) staining was scored, regardless of staining intensity, in all endometrial cancer glands contained within triplicate repeat cores.
